# A predictive model of fractional land use

**DOI:** 10.1101/2020.12.08.415992

**Authors:** Simon Kapitza, Nick Golding, Brendan A. Wintle

**Affiliations:** School of Biosciences, The University of Melbourne, Parkville VIC 3010, Australia; Telethon Kids Institute, Perth Children’s Hospital, 15 Hospital Ave, Nedlands WA 6009, Australia; Curtin University, Kent St, Bentley WA 6102, Australia

**Keywords:** Land use forecasting, fractional land cover, continuous fields, agricultural expansion, socio-economic change, biodiversity conservation, cross-validation, multinomial regression

## Abstract

1. Land use change leads to shifts in species ranges and declines in biodiversity across the world. By mapping likely future land use under projections of socio-economic change, these ecological changes can be predicted to inform conservation decision-making.
2. We present a land use modelling approach that enables ecologists to map changes in land use under various socio-economic scenarios at fine spatial resolutions. Its predictions can be used as a direct input to virtually all existing spatially-explicit ecological models.
3. The most commonly used land use modelling approaches provide binary predictions of land use. However, continuous representations of land use have been shown to improve ecological models. Our approach maps the fractional cover of land use within each grid cell, providing higher information content than discrete classes at the same spatial resolution.
4. When parametrized using data from 1990, the method accurately reproduced land use patterns observed in the Amazon from 1990 until 2018. Predictions were accurate in terms of the fractional amounts allocated across the landscape and the correct identification of areas with declines and increases in different land uses. A small case study showcases the successful application of our model to reproduce patterns of agricultural expansion and habitat decline.
5. The model source code is provided as an open-source R package, making this new, open method available to ecologists to bridge the gap between socio-economic, land use and biodiversity modelling.

## Introduction

Land use change is a key driver of global environmental change, causing global declines in biodiversity, species extinctions and resulting in the deterioration of ecosystem ser-vices (Foley, 2005; IPBES, 2019).

Land use change is driven by bio-physical and socio-economic processes (Lambin et al., 2011). Climate change will likely result in global shifts and declines of land suitable for agricultural production, with projected depletion of land reserves in the first half of the 21st century (Lambin et al., 2011). Most socio-economic scenarios of future change describe future increases in food production and international trade of goods (O’Neill et al., 2014, 2017). Even under lowest impact scenarios, also known as ‘shared socio-economic pathways’ (SSPs), in which land use is strongly regulated, deforestation rates are reduced, diets are more plant-based and climate change mitigation starts early, crop and livestock production are still likely to be higher and occupy a larger land area than they do today.

Despite mounting evidence of adverse environmental impacts of historic and current land use change, work concerned with understanding future biodiversity change tends to focus on climate change (Titeux et al., 2016; Struebig et al., 2015), or other aggregated effects of socio-economic change, such as forest loss (Margono et al., 2014) and urban expansion (Seto et al., 2012). Consequently, future predictions of biodiversity change will benefit from explicit accounting of the drivers and effects of land use change at the level of individual types of use. Detailed, large-scale mappings of future land use will provide invaluable insights for researchers and policy makers, particularly in terms of conservation planning and preventing future biodiversity loss.

Different conceptual approaches have found application in investigations of past, present and future land use change. van Schrojenstein Lantman et al. (2011) identify four theoretical core principles for modelling land use change. Predictions can be made based on *the continuation of historical developments*, where past patterns are extrapolated into future conditions, the *suitability of land*, where land use changes are predicted based on proximity to markets, biophysical conditions and other environmental drivers, *neighbourhood interactions*, where neighbouring land uses affect local changes and *actor interactions*, where land use changes explicitly emerge from the decision-making of individual actors, or groups of actors.

These principals appear in many existing models. For example, artificial neural networks and markov chain models learn and infer patterns from historic time series of land use change (Tayyebi and Pijanowski, 2014; Pijanowski et al., 2002). To allow for spatially-explicit assessments, markov chain models have been frequently combined with cellular automata (CA-Markov models, see Hyandye and Martz, 2017; Aburas et al., 2017; van Schrojenstein Lantman et al., 2011). In cellular automata the transition probability of a cell to another land use depends on its current state and the state of neighbouring cells, both of which are the result of historic changes (van Schrojenstein Lantman et al., 2011). Cellular automata have been used successfully to simulate strongly auto-correlated changes, such urban sprawl (Verburg et al., 2004b; Fang et al., 2005; Shafizadeh Moghadam and Helbich, 2013; Sun et al., 2007).

Some existing modelling approaches apply regression analysis and other techniques to identify associations between various environmental conditions and observed land-use patterns (Meiyappan et al., 2014; van Schrojenstein Lantman et al., 2011; Lambin et al., 2000; Verburg et al., 2004b). Some of the most prominent examples include the Conversion of Land Use and its Effects (CLUE) model series, which have found application in the prediction of spatially-explicit patterns of land use at national and continental scales (Veldkamp and Fresco, 1996b; Verburg and Overmars, 2009; Verburg et al., 1999, 2002; Kapitza et al., 2020). Exogenously determined future changes in area demands for different land uses, often predicted by an economic model (Aguiar et al., 2016), may be downscaled by establishing statistical relationships between observed land use and a set of socio-economic and bio-physical drivers of land use and land use change. Predicted land use suitability surfaces inform local competition for different land uses (Verburg et al., 2002; Meiyappan et al., 2014). Models can be further parametrized by including transition rules at local (cell) and landscape levels and constraints on overall turn-over through time. More simplistic models based on statistical analysis use an ordered allocation algorithm, in which competition between land uses is handled by ordering allocations in terms of perceived socio-economic value (Fuchs et al., 2013).

Modelling approaches based on statistical analysis are useful in particular because of their transparency and scalability from regional to continental levels. For example, Veldkamp and Fresco (1996a) show that relationships between land use and biophysical and human driving factors in Costa Rica act differently at different scales, high-lighting the importance of the model’s capability to parametetrize relationships in close consideration of the study area.

Most land use models apply statistical analyses of discrete land use classes using binary logistic regression to model the cell-wise probabilities of occurrence for each land use, independent of the probabilities of other land uses. The resulting probability of land use occurrence at a site produced by separate models is an incomplete representation of the underlying structure of land use probability, because it omits that occurrence probabilities are dependent between land use types, and that the probabilities of all discrete classes must sum to one. For example, when a site has very high probability for urban land use, this implies relatively low probabilities for primary natural habitat, which separate, independent logistic regressions do not fully capture.

One step toward explicitely modelling competition between land uses is to apply multinomial regression, thus allowing for the prediction of conditional binary probabilities of multiple classes (Noszczyk, 2019). However, the classifier would still allocate the land use with the highest probability at a site. For many ecological considerations it is desirable to know individual probabilities of land use occurrence for each land use type in order to characterize the underlying continuous fractions occupied by different land use types within a classified site. A few model examples are capable of predicting continuous fractions of land use at very coarse resolutions (see Hasegawa et al., 2017; Meiyappan et al., 2014), but documented approaches are not yet available in a usable package suited to regional-continental scale.

Increasingly, categorical data sets are available at spatial resolutions of finer than 1km^2^. Three prominent examples include the CORINE Land Cover inventory (Bossard et al., 2000), which contains several time steps between 1990 and 2019 at 100m resolution for the European continent, global land cover mappings produced for the year 2010 through Copernicus Land Monitoring Service (European Union, 2019) at the same resolution, as well as global mappings of land cover in annual time steps between 1992 and 2018, produced under the European Space Agency’s (ESA) Climate Change Initiative Land Cover (CCI-LC) project (ESA, 2019), available at 300m resolution.

However, the spatial variables that represent drivers of land use and biodiversity change are often not available over large spatial extents at fine resolutions above 1km^2^ (Dendoncker et al., 2006), though this situation is changing. Global mapping and climatic projections based on Global Circulation Models (Hijmans et al., 2005) and other drivers, such as soil properties (Global Soil Data Task Group, 2000), are all now mapped at 1km^2^ or better. Infrastructure such as roads (Center for International Earth Science Information Network - CIESIN - Columbia, 2013) and built-up areas (FAO, 1997) is typically represented by geographic features, but can be converted to raster representation at fine resolutions.

Lowering the resolution of available land use data sets to fit the resolution of continental- or global-scale environmental covariates has the advantage of higher computational efficiency when simulating changes. Assigning a single category of land use on each larger pixel effectively eliminates sub-pixel information on land use (Seo et al., 2016), so this approach is not desirable. In order to retain information, it is preferable to calculate the fractions of land use covering each new cell, producing continuous fields of information and retaining information at sub-pixel level (Seo et al., 2016).

Mapped representations of land use fractions have high utility in down-stream ecological modelling applications because they preserve information on spatial heterogeneity within classes, thus providing a much more refined landscape representation. For example, many wide-ranging species may persist in landscapes if a certain proportion of the landscape is comprised of old forest. It has been shown that continuous fields of land use allow better estimation of biomass and biomass change (Xian et al., 2015) and are better able to explain variation in home range sizes (Bevanda et al., 2014) than categorical land use data. Continental-scale biodiversity assessments have shown that patterns are associated with high spatial-resolution fractional land use measures such as the regional aggregation of land use types, land cover diversity and land use covariates including land use intensity (Mouchet et al., 2015) and actual evapo-transpiration (Mouchet et al., 2015; Whittaker et al., 2006). Creating mappings of some of these co-variates requires fine-scale mappings of fractional land use as principal input (Plutzar et al., 2016). The intensification of agriculture and forest harvesting are crucial factors shaping biodiversity (Levers et al., 2014, 2016) that require inputs of crop type and vegetation composition within each spatial unit. These ecological considerations of the utility of fractional land cover and land use representations are underpinned by recent advancements in algorithms to produce high resolution mappings of fractional land cover from satellite data (Allred et al., 2020; Hill and Guerschman, 2020).

Here, we innovate a land use modelling approach that allows ecologists to incorporate fractional land use change into ecological modelling. An advantage of our approach compared to existing fractional land use modelling approaches is its ease of of implementation for ecologists trained in R (R Development Core Team, 2008) and its high scalability to high resolutions and large spatial extents with minimal parametrization requirements. The source code for our method is freely available as a small open source R package hosted on GitHub (https://github.com/kapitzas/flutes). As such, our approach contributes a new open method toward bridging the gap between socio-economic, land use and biodiversity modelling.

We provides a mathematical description of the developed fractional land use model and evaluate our model according to its ability to correctly estimate the direction and intensity of observed land use changes using a case study in the Brazilian Amazon.

## Materials and methods

### Model description

The model consists of two main components (Fig. 1). First, statistical analysis is used to determine how the suitability of the landscape for different land uses relates to a set of environmental drivers of land use change, producing a suitability surface for each land use class (Fig. 1a). Second, fractional changes in additional land use demands are allocated iteratively in the landscape, scaling with the land use suitability surfaces (Fig. 1b). We utilize a cellular automaton to introduce cell-level allocation decisions that constrain the location and direction of land use changes according to three rules:

**Figure 1:**
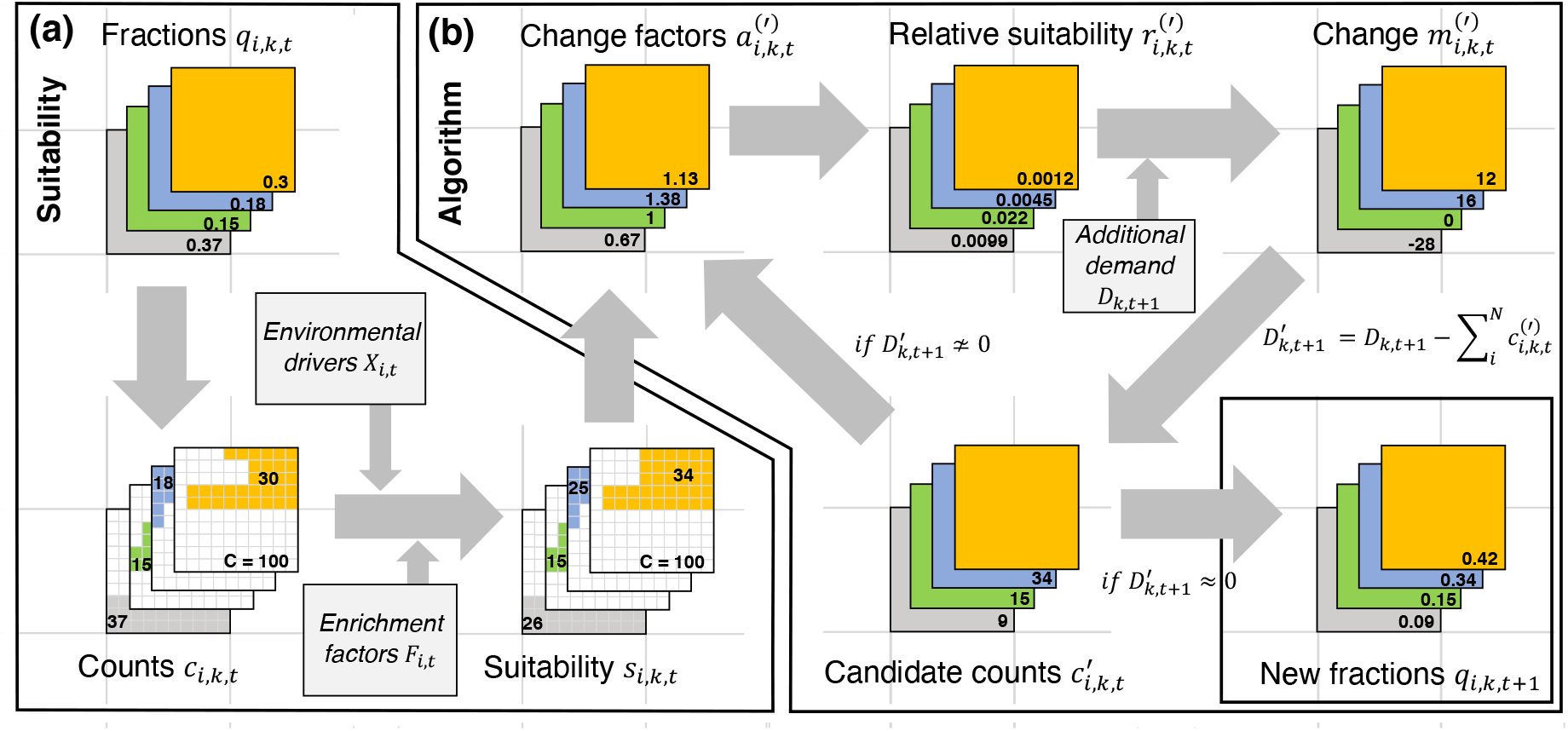
Conceptual diagram of land use modelling approach. a) Land use suitability model. Observed fractions of land use are first converted to integer counts through multinomial draws and their relationship with environmental drivers and neighbourhood covariates (derived from previous time step’s land-use distribution) is assessed. b) Allocation algorithm. First it is estimated by how much each cell has to change to achieve the modelled ideal distribution of land uses. Change factors are then converted to relative suitabilities that serve to distribute land use supply required to satisfy the additional demand in the landscape. Multinomial draws assure that each cell’s land use class probabilities sum to 1. The resulting difference of the current supply and the total additional demand is recalculated to support allocation in the next iteration. The cycle repeats until the difference between the current supply and total additional demand is very close to zero, meaning that all additional demand has been allocated. At this point, the integer counts representing the land use fractions on each cell are converted back to fractional representation

#### Rule 1: Future land use supply meets additional demand

Projections of land use demands may be provided through external models, such as Computational General Equilibrium (CGE) models (i.e. GTAP, Aguiar et al. (2016)), or through the analysis and extrapolation of historic patterns (Moulds et al., 2015). The model allocates additional demand by adding cell-level supply *d*_*i,k,t*+1_ in cell *i*, land use *k* and time step *t* + 1 to fractions *q*_*i*,*k*,*t*_ (Fig. 1b). The first model objective can be formulated:

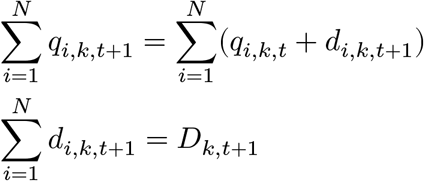

*D*_*k,t*+1_ is the additional landscape-wide supply and is at equilibrium with additional demand after the algorithm converges.

#### Rule 2: Cell-level fractions must sum to 1

The second model condition requires that supply *d*_*i,k,t*+1_ is allocated across cells in such a way that 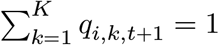 (Fig. 1b).

#### Rule 3: Allocations are determined by land use suitability

The third model condition requires cell-level supply *d*_*i*,*k*,*t*+1_ to be distributed in such a way that the allocated amounts in each cell scale with a predicted probability surface *s*, by modelling 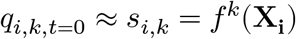, where *X*_*i*_ is a set of demographic and bio-physical drivers related to land use. *f*^*k*^ is a multinomial, multi-response model (Fig. 1a). The parameter estimation of this model is based on the first time step and predicted to the conditions of subsequent time steps.

The land use status in a cell’s neighbourhood has been shown to play an important role in determining a cell’s land use (Dendoncker et al., 2007; Mustafa et al., 2018; van Vliet et al., 2013; Verburg et al., 2004a). Our suitability model applies neighbourhood interactions by calculating autocovariates (Verburg et al., 2004a) and including these in the multinomial regression of the land use suitability model. Following Verburg et al. (2004a), our autocovariates measure the amount of clustering of land uses in the cell neighbourhood when compared to the entire landscape. We calculate autocovariates as enrichment factors 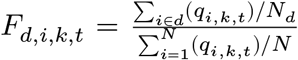. The numerator is the average fraction the average fraction of land use *k* in the neighbourhood *d* of each central cell *i* and the denominator is the average fraction of land use *k* in the entire landscape *N*. Here, we only included neighbourhood characteristics in the 3×3 neighbourhood around each central cell, but other neighbourhoods are possible (Verburg et al., 2004a). When predicting suitability at each time step, the autocovariates are recalculated based on the assigned fractions from the previous timestep.

The response here is represented by fractional land use and not discrete classes normally required in multinomial regression. Therefore, we assume that underlying the land use fractions for each cell is a vector of counts *c*_*i,k,t*_ that sums to a total number of counts *C* in each cell (e.g. *C* = 1*e*6). We derive these counts through *c*_*i*,*k*,*t*_ ≈ *q*_*i*,*k*,*t*_ * *C*. In integer representation, the data are approximately proportional to the original fractions. When fitting the suitability model, parameter uncertainty depends on the assumption of *C*. *C* should be chosen to represent the degree of numerical precision in the observed fractions. I.e. if there are only 2 decimal places, setting *C* = 100 results in counts that represent all of the information contained in the original fractions. Accordingly, the multinomial logit model takes the form

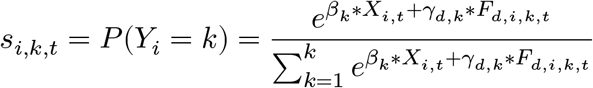

where *k* is the reference land use class, *β*_*k*_ the estimated parameters in each class for covariates *X*_*i,t*_ and *γ*_*d,k*_ the estimated parameters for autocovariates *F*_*d*,*i*,*k*,*t*_. We estimated parameters using R’s ‘nnet’ package (Venables and Ripley, 2002). Predicted fractions satisfy 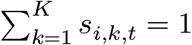.

All software development and model validation was conducted in R (version 4.0.1) (R Development Core Team, 2008).

### Data

We developed and tested our model using land use and environmental data from the Amazon basin. We downloaded and cropped 7 time steps (1992, 1997, 2003, 2008, 2013, 2015 and 2018) of the global land cover mapping provided through the European Space Agency’s Climate Change Initiative Land Cover (CCI-LC) project (ESA, 2019). These data are available at a grid resolution of 300m. We combined the recorded 31 land cover classes to 9 new classes of land use we deemed crucial to identify processes leading to agricultural expansion and declines in habitat (Table 1). We aggregated the resolution to 10km^2^, calculating fractions of land use from the cell counts of each land use class on the original map present in each new cell. Fractional land use in *K* classes is mapped over *N* raste1r cells, with fractions *q*_*i*,*k*,*t*_ in cell *i* in each land use class *k* always satisfying 0 ≤ *q*_*i*,*k*,*t*_ ≤ 1 and 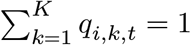.

**Table 1:** Mapping of original land use classes to new classes applied in this study

|  | New class | Abbr. | CCI-LC class | Description |
| --- | --- | --- | --- | --- |
| 1 | Cropland | Cro | 10, 11, 12, 20, 30 | Rainfed and irrigated cropland, mosaic cropland with >50% cropland and natural vegetation (tree, shrub, grass) |
| 2 | Cropland mosaic | CrM | 40 | Mosaic cropland with <50% cropland and natural vegetation (tree, shrub, grass) |
| 3 | Forest | For | 50, 60-62, 70, 80, 90, 100, 160, 170 | Forest, closed to open, with >15% canopy cover, Mosaic tree/shrub (>50%) / herbacious cover, Flooded tree cover |
| 4 | Grassland | Gra | 110, 130 | Grassland and mosaic herbacious cover (>50%) / tree/shrub |
| 5 | Shrubland | Shr | 180 | Flooded shrub or herbacious cover |
| 6 | Wetland | Wet | 190 | Settlement, Urban land uses |
| 7 | Urban | Urb | 120 | Closed to open and open shrubland |
| 8 | Other | Oth | 140, 150, 151-153, 200-202, 220 | Lichen/mosses, sparse trees/shrubs/herbaceous vegetation, bare areas, snow/ice |
| 9 | Inland water | Wat | 210 | Natural and artificial inland water bodies |

We downloaded a set of spatially explicit climate, topographic soil and human covariates (Table 2 for a full list of covariates), derived neighbourhood covariates from observed land use in the first time step and estimated observed demand change by calculating the landscape-wide mean fraction for each land use class in each observed time step. All explanatory covariates were standardized to have mean 0 and standard deviation 1. We removed covariates from correlated pairs (Spearman’s rank correlation coefficient > 0.7), always retaining the covariate with the smaller average correlation with all other covariates in order to maximise the amount of independent information in the final data set used for fitting.

**Table 2:**
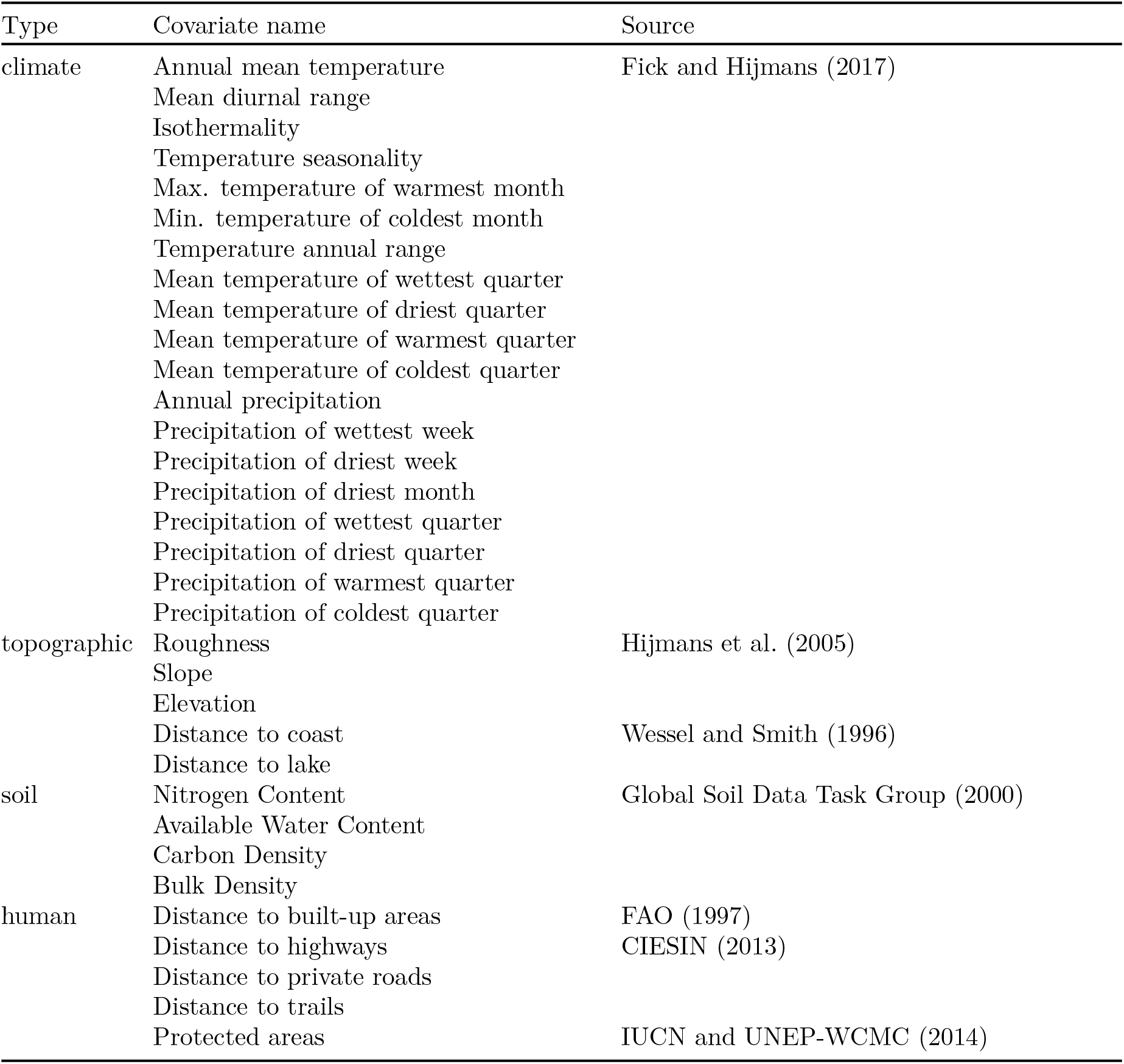
List of covariates that were included in land use suitability model

### Model constraints

Analysing time series data, we determined that only very small percentages of cells change from being devoid of a particular land use to containing that land use within one time step (Table 3). Therefore, we added a constraint that land use increases are more likely to be applied to cells where the land use is already present. The constraint parameter was the percentage of cells in which a non-existent land use was newly established between time steps. For example, setting the constraint to 100% would allow increases of a land use in all cells that did not contain that land use in the previous time step.

**Table 3:**
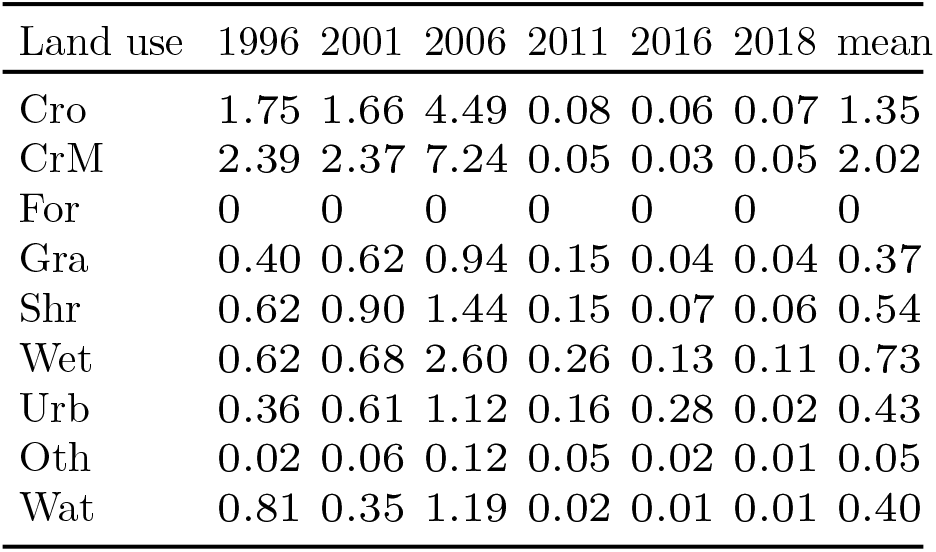
Share of cells (%) containing a land use that were completely void of that land use in preceding time step. Values derived from observed time series.

We parametrized the constraint by determining the time series mean of the according percentages between all time steps for each class (Table 3). For example, throughout the simulation, we allowed *Cro* increases in 1.35% of the cells in which *Cro* was not present in the preceding time step (Table 3). From cells currently zero in a land use, we selected the ones for increases that had the highest predicted land use suitability for that land use.

We masked category I and II protected areas established up until 1992 from land use changes as has been shown previously (see Fig. 2 for a map of protected areas) (Verburg et al., 2002; IUCN and UNEP-WCMC, 2014; Kapitza et al., 2020).

**Figure 2:**
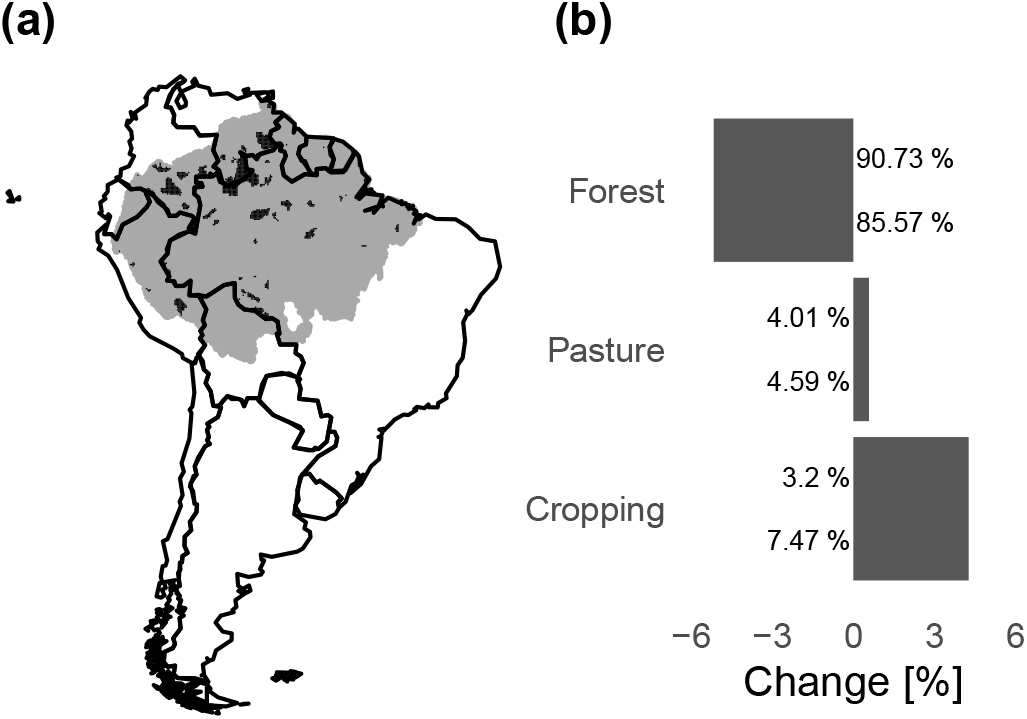
Overview of the study area. a) Location of the amazon catchment in South America (grey-shaded area), including IUCN protected areas (categories I and II) which were used to constrain land use changes (black shaded areas). b) Changes in selected land uses, derived from observed land use maps. Pasture includes Gra and Shr, Cropping includes Cro and CrM and forest includes For. Land use classes are specified in Table 1 below

### Validating the intensity and direction of predicted changes

First, we examined the accuracy of the multinomial suitability model and how it is affected by spatial resolution and the included covariates. To account for spatial autocorrelation in the environmental covariates and land use time series, we conducted spatial-blocks cross-validation (Valavi et al., 2019) by separating the landscape into 9 equal-sized spatial blocks. We fitted models using data from 8 of the 9 blocks and predicted the model to the withheld block, until predictions were made for the entire study area. We cross-validated suitability models at 1km^2^ and 10km^2^, including 1) only environmental covariates, 2) only neighbourhood covariates and 3) both co-variate types combined. We conducted correlation analysis and removed highly correlated covariates from pairs, always keeping the covariate with the lower average correlation with all other covariates in order to maximise the amount of independent information retained in the covariate set. For each of the three models we measured predictive performance by estimating cell-level Suitability Root Mean Squared Error (RMSE_suit_) between the suitability surfaces *s*_*m*,*i*,*k*,*t*_ and the observed fractions *o*_*i*,*k*,*t*_, following 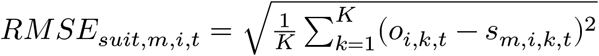 for each suitability model *m*.

Second, to validate the intensity of changes predicted by the allocation algorithm, we assessed the accuracy of predictions of cell-level fractions under a null model, a naive model, a semi-naive model and a fully parametrised model throughout the observed time series. 1) Under the null model, we assumed no change of land use through time. The null model served as reference to measure the improvements provided by each additional model component. 2) Under the naive model we only allocated additional demands, but scaled cell-level allocations with the average supply observed across the entire landscape. This model assumes that suitability is not informative about where a change will happen and that allocations are equally likely to be anywhere in the landscape. 3) Under the semi-naive model, cell-level allocations were additionally scaled with the predicted suitability surfaces *s*_*i*,*k*,*t*_ (as illustrated in 1). 4) Under the full model, allocations were scaled with suitability surfaces *s*_*i*,*k*,*t*_ and all constraints (constraining most increases to cells where land use type already exists and masking protected areas from changes) were applied.

We calculated RMSE_alloc_ under each allocation model to estimate how well the different model components simulated each cell-level vector of land use fractions *q*_*m*,*i*,*k*,*t*_ compared to the respective observed vectors *o*_*i*,*k*,*t*_, following 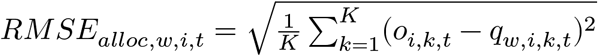.

Due to the squared term, RMSE cannot inform on whether the models correctly identified the direction of change. Therefore, we estimated and validated the direction of cell-level changes (decreases, no change, increases) separately. We mapped these transitions for each class between the time steps of the observed time series and the time steps of the time series simulated under each model. We calculated *overall difference* of each pair of corresponding maps to obtain an interpretable measure of similarity of predicted and observed direction of changes (Pontius and Millones, 2011; Pontius and Santacruz, 2014). Achieving high accuracy in these first two model goals would suggest that simulated patterns of land use change closely resemble observed patterns.

### Case study: agricultural expansion in the Amazon Basin

The Amazon catchment is largest river basin in the world and occupies over one third of the South American land mass (Fig. 2a). As the world’s most diverse tropical forest area, the basin hosts at least 10% of the world’s known species (Da Silva et al., 2005).

The Amazon biome is threatened by a multitude of interacting factors. Ecosystem services, such as water supply, carbon storage and provision of species habitat are directly threatened by the effects of climate change and the increasing pressure on land, with projected severe reductions in water yields, carbon content and species habitat, which is particularly affected by changes in natural vegetation cover (Prüssmann et al., 2016). The primary uses for cleared forest land are pasture for cattle farming and industrial soy cropping (Nepstad et al., 2014; FAO, 2015). Between 1992 and 2018, the biome has seen significant increases in land required for cropping and pasture, as well as significant decreases in forest cover (Fig. 2b).

Using a broad reclassification of the predicted and observed land use classes into crop-land, pasture and habitat, we were able to specifically validate our model’s ability to predict agricultural expansion and habitat declines as aggregated threats to ecosystems and biodiversity. We assessed the accuracy of our predictions of cropland expansion with simultaneous declines in classes containing natural habitats (*For*, *Wet* and *Oth*). We categorized the observed and predicted maps into 1) areas with no cropland increase, 2) areas where cropland increase led to mostly forest declines (net replacement of forest), and 3) areas where cropland increase led to mostly declines in other natural habitat classes (net replacement of other habitat). Similarly, we assessed the accuracy of our predictions of pasture expansion on natural habitats by categorizing the landscape into 1) areas with no pasture increase, 2) areas where pasture increase led to mostly forest declines, and 3) areas where pasture increase led to mostly declines in other natural habitat classes. We assessed the difference between the respective observed and predicted maps by disaggregating overall difference into allocation and quantity difference components. This allowed us to further investigate whether prediction inaccuracies were due to error in the sum of allocated cells (quantity difference), or to errors in spatial location, discounting quantity difference (allocation difference) (Pontius and Millones, 2011).

## Results

### Predicting land use change intensity

Results of the cross-validation of the suitability model component show that including neighbourhood covariates resulted in substantial predictive performance improvements across spatial blocks (Fig. 3c) at both resolutions; models using neighbourhood co-variates alone were approximately as good as the model using the full covariate set. Including only environmental variables resulted in less accurate predictions at both resolutions, with predictions under the fine resolution comparatively worse than under the coarse resolution.

**Figure 3:**
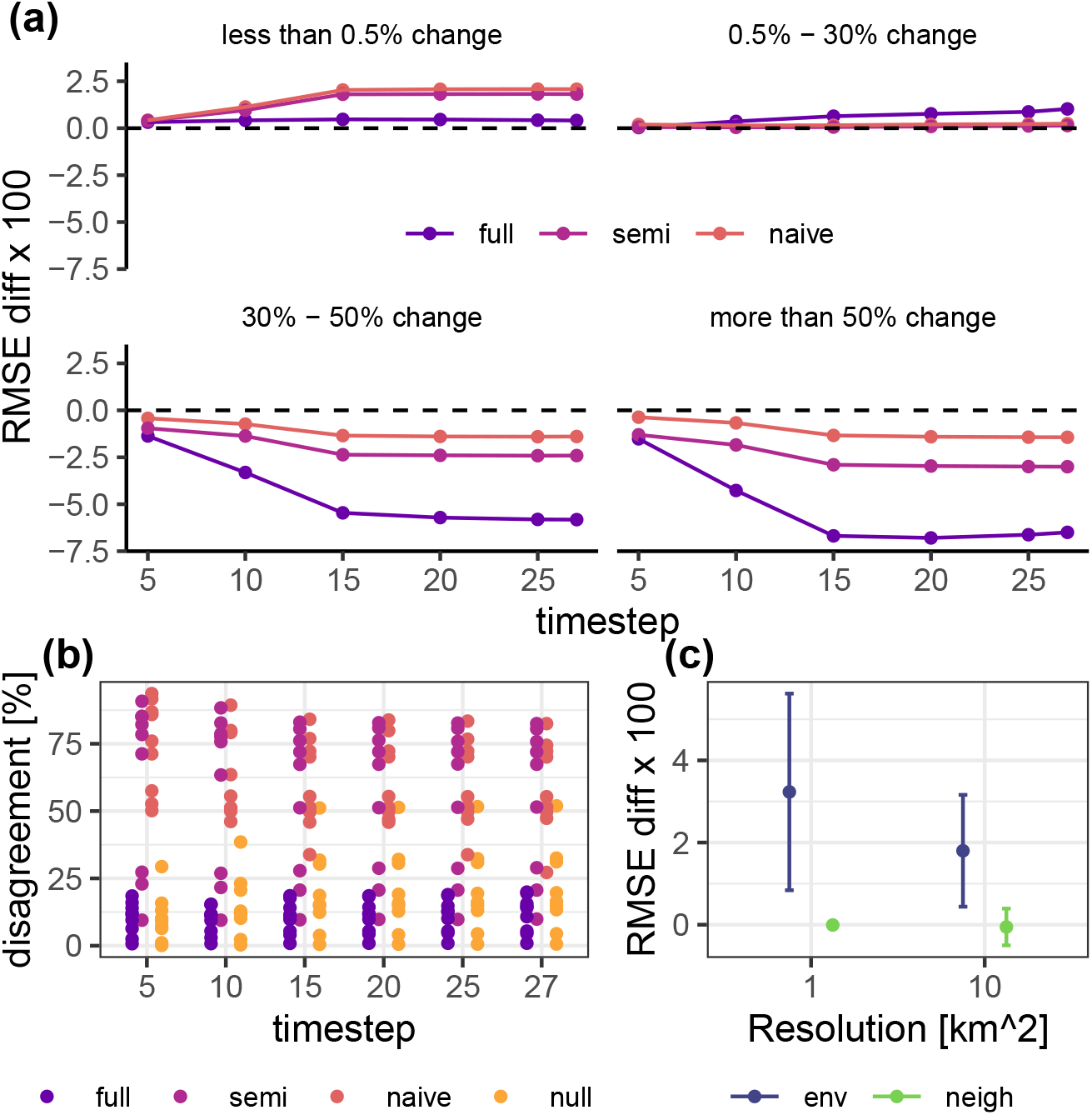
Validation of predicted land use change intensity and direction of change and cross-validation of suitability model. a) The difference between RMSE for each model (naive, semi-naive, full) and RMSE of the null model. The null model assumes that land use is static through time, the naive model assumes completely random allocations, the semi-naive model assumes that allocations are scaled with land use suitability and the full model assumes that allocations are both scaled with land use suitability and subject to model constraints (no changes in areas under high protection status and no land use increases in areas completely devoid of that land use). All RMSE were calculated at cell-level, using the predicted and observed vectors of land use fractions in each cell. Plotted are means and standard deviations across cells. Positive values indicate better fits under the null model, negative values indicate better fit under more highly parametrised models. Data on validation outcomes are grouped by the magnitude of the largest observed proportional change in any land use within a cell. In general, the larger the observed change in land use, the better the parameterized models did compared with the null model. b) The proportional disagreement between predictions of the direction of change (no change, decrease, increase) for each land use and the observed direction of change at each time step. Smaller values indicate lower overall difference and higher similarity between corresponding maps. c) Difference between cross-validated RMSE estimated for suitability models containing only environmental covariates and only neighbourhood covariates and models containing both covariate types combined. Positive values indicate a poorer fit than the model containing both covariate types.

Under all tested models (naive, semi-naive, full), the accuracy of cell-level allocations improved with the intensity of observed changes (Fig. 3a). This implies that our model makes good predictions under scenarios with high expected overall changes. It is not surprising that large changes are easier to predict than small ones.

Where observed changes were large (Fig. 3a, bottom two panels), including land use suitability and constraints (full model) resulted in substantial increases of predictive performance. In these areas, the null model’s assumption of no spatial variation in reallocation of land use introduced very high bias, which our constraints were able to reduce.

When observed changes were small (Fig. 3a, top two panels), the null model made near perfect predictions. Given how close the null model already was to the truth, improvements by allocating demand (naive model) and accounting for land use suitability (semi-naive model) were difficult to achieve; in the smallest change category (Fig. 3a, top left panel), the naive and semi-naive predictions were in fact slightly worse than the null. In these areas the largest observed changes were below 0.5%, making the assumption of no change under the null model highly plausible. Under the full model, the applied constraint limited the areas that could be flagged for increases. Accordingly, where observed changes were small, this model made better predictions than the semi-naive and naive models, in which this constraint was not applied.

### Predicting the direction of land use changes

The worst predictions of cell-level direction of change were made by the naive and semi-naive models and the best predictions under the full model (Fig. 3b), with overall difference consistently less than 25%. Predictions became more accurate the more model components were applied. Under the full model we achieved the highest prediction accuracy. Overall, the semi-naive model performed slightly better than the naive model, demonstrating the utility of scaling allocations with land use suitability surfaces. However, both naive and semi-naive predictions of change were generally less accurate than those under the full model. For areas with very small changes predictions tended to be worse than those under the null model. This was due to the large number of cells falling into the smallest category of observed change (≈ 60% of cells when measured across the entire time series); in such areas, assuming the null model was more accurate than making naive and semi-informed allocations. In areas with intermediate levels of change (0.5%-30%), predictions under the different models were very similar, but the range of estimated RMSE was higher under the full model (indicated by wider error bars). This suggests that the full model outperformed the semi-naive and naive models in some areas, but fell comparatively short in others. This may be due to a larger number of cells at this level of observed change incorrectly prevented from increasing in land uses by the applied model constraints, while similarly a larger number of cells was correctly allowed to increase by the model.

### Predicting agricultural expansion and habitat declines

Our model achieved high accuracy when predicting cropland and pasture expansion on forest and other land use types containing natural habitats. The estimated overall difference between observed and predicted mappings was consistently around 9% when measuring the impacts of cropland expansion (Fig. 4a) and 16-18% when measuring the impacts of pasture expansion (Fig. 4b).

**Figure 4:**
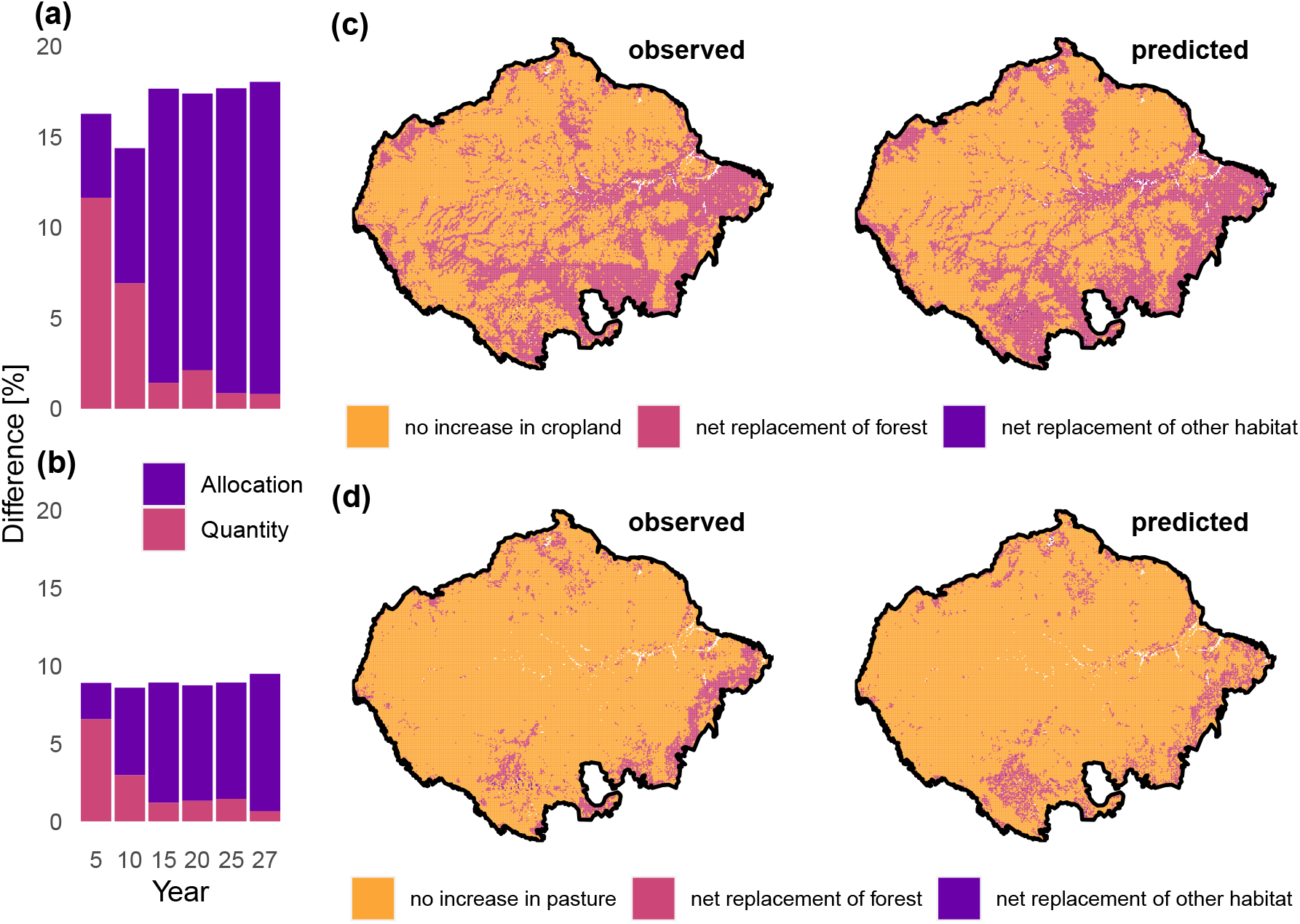
Validation of predictions of aggregated agricultural expansion and natural habitat decline in the Amazon basin. a,b) Quantity and allocation difference of observed and predicted mappings of agricultural expansion (a) and natural habitat decline (b), by time step. The overall difference is the sum of quantity and allocation difference. c) Maps of observed (left) and predicted (right) cropland expansion and habitat declines between 1992 and 2018. d) Map of observed (left) and predicted (right) pasture expansion and habitat declines. Quantity and allocation difference on these maps correspond to the respective last bars in panels a and b.

Comparing the spatial configuration of cropland expansion (Fig. 4c) and pasture expansion (Fig. 4d) into natural habitats in the last validation time step, the number of cells falling into each category was very similar in both cases, but the spatial arrangement differed, illustrating the different relative contributions of quantity and allocation difference. The model overestimated cropland expansion and net forest loss in the central-north, east and south of the catchment, while small observed areas in the center of the catchment were slightly underpredicted. Cropland expansion leading to net losses in non-forest habitats were barely visible in both the observed and predicted maps. Pasture expansion leading to net forest loss was slightly underestimated in the north and east of the catchment, and slightly overestimated in the center-south. Pasture expansion leading to net losses in other habitat types was very small.

## Discussion

We have presented a new land use model to predict land use fractions, thus retaining information at sub-pixel level. The model is able to accurately allocate fractions of land use through time, especially under scenarios of more extreme land use change. We explicitly accounted for competition between land use types and land use suitability in response to environmental drivers by means of a multinomial logistic model and could show that this aspect brings substantial improvements to predictions, when compared the assumption that land use does not change at all (null model).

In scenarios where demand changes are expected to be high, our model allocates supply to match aggregated demand, changing the total area allocated to different land uses and also allowing land uses to be established in new areas. In scenarios with small expected demand changes, land use changes, including the establishment of land uses in new areas, remain small.

The initial land use distribution is likely to have resulted from long time periods of optimizing behaviour. For this reason, our model assumes that the land use distribution does not change to match predicted land use suitability alone. For example, if the modelled cropland suitability in an area is 0.8, but the observed cropland fraction is 0.2, there would only be a local increase in cropland if the aggregated demand for cropland at the study area level increased. The much lower realised fraction of cropland in that area when compared to the predicted suitability for cropland captures processes that are not captured by the suitability model.

Similar to CLUE, our constraint on turn-over accounted for conversion effort. Here, data from the observed validation time series allowed us to extract a raw estimate of the constraint parameter to tune our model. We estimated the parameter using long-term observed means, which we assume to be similarly informative as extensive literature review, inquiring expert opinion, or analyzing data from time series preceding the predicted time span, thus preventing overfitting.

We could show that our model is very easily adaptable to specific ecological study contexts. When validating our model’s performance in the context of agricultural expansion on natural habitat, we divided the difference between predicted and observed maps of agricultural expansion and habitat decline into the two components quantity and allocation difference. We were able to determine that the main sources of difference between predicted and observed maps were spatial misallocations that increased with increasing time horizon. However, allocation difference was still very low, suggesting that our model can be a useful tool to predict the overall pressure and spatial configuration of land use change impacts that are driven by different types of agricultural expansion into different habitat types.

Validating the suitability model component of our model approach, we found that neighbourhood covariates explained much of the suitability patterns across the land-scape. This is a common effect of including flexible spatial correlation terms in models with other spatially-varying covariates (spatial confounding) (Hodges and Reich, 2010). The models describe the spatial pattern with the spatial correlation term, but this effect does not imply causation and other drivers included in the model may still drive changes in the response, particularly over long time periods. Here, similar to what was shown by Dendoncker et al. (2007), including neighbourhood covariates lead to the most highly fitted models. Allowing spatial autocorrelation to drive patterns seems a sensible choice for predictions in this case study because the model only predicts three decades. How-ever, for longer time spans, spatial autocorrelation probably becomes less important and continental-scale environmental driving factors acting homogeneously across the whole landscape may dominate patterns in reality. When making such longer-term pre-dictions, this could be captured by fitting the suitability model with several time steps of data, thus assuring that land use suitability is less reliant on the present land use state, but more weight is given to long-term and large-scale environmental processes.

The results of our validation also strongly indicate that in case of our model, adding constraints (decision rules) in terms of where and how land use changes are allowed to occur, are responsible for the majority of increases in predictive performance. While we provide initial steps in parametrising these constraints, more specific knowledge of bottom-up processes that drive land use stasis and change across the landscape could further consolidate the accuracy of our model. For example, this could be achieved by including data on the expected behaviour of economic agents who seek to maximise returns on their productive land. One example includes the Land Use Trade-offs (LUTO) model (Bryan et al., 2014; Connor et al., 2015), which includes pixel-wise optimisation of cost and return of alternative land uses. However, such models are difficult to parametrise in data-scarce regions and require significant computational power. Bottom-up processes, such as price feedbacks, also tend to act at very fine spatial resolutions, but have little effect when seen at a continental scale, where scenario uncertainty and global processes dominate predictions (Connor et al., 2015). Depending on scale, including very fine-scale dynamics of agent behaviour may simply not pay off, or it might be more appropriate to merely downscale them to the study area extent (Van Asselen and Verburg, 2013; Connor et al., 2015).

In order to allow scaling our model to global applications, we only used drivers that were available at global scales. However, improvements to the land use suitability model can be achieved by including more proximate drivers of land use change, such as market accessibility (Meiyappan et al., 2014; Verburg et al., 2011), by fitting the land use suitability model for individual subsets of the study area to improve local fit, or by creating more land use classes for which particular biophysical constraints are known. Including location-dependent drivers and models and raising the resolution may substantially improve the accuracy of land use suitability maps, increasing the contribution of this model component to overall prediction accuracy.

Our approach provides a validated method to spatially downscale future changes in land demands, and we highlight options to further improve its applicability in ecological studies. We hope that by providing open source code we can encourage ecologists to include land use change in predictive studies and make further steps toward consolidating quantitative methodological links between socio-economic and ecological systems.

## Acknowledgements

This work received funding under the Australian Research Council Discovery grant DP170104795. NG was supported by an ARC DECRA fellowship (DE180100635). The authors acknowledge contributions made throughout the research phase by J. Elith and D. Zurell. SK was supported by the Melbourne International Research Scholarship (MIRS).

## Author contributions

SK, BW conceived the idea of providing a fractional land use model. NG, SK and BW designed the model and validation. SK coded the model and analyzed the data. SK led the manuscript with edits from NG and BW.

## Data availability

All inputs required to repeat this work will be made available through a FigShare repository upon publication. An R package containing the model source code is available on GitHub (https://github.com/kapitzas/flutes). All code for data preprocessing and analysis is also provided through a GitHub repository (https://github.com/kapitzas/frac_lumodel). Both repositories will be receive permanent DOI through Zenodo upon publication.

